# Recapitulation and Retrospective Prediction of Biomedical Associations Using Temporally-enabled Word Embeddings

**DOI:** 10.1101/627513

**Authors:** Jiho Park, Agustin Lopez Marquez, Arjun Puranik, Ajit Rajasekharan, Murali Aravamudan, Enrique Garcia-Rivera

## Abstract

The recent explosion of biomedical knowledge presents both a major opportunity and challenge for scientists tackling complex problems in healthcare. Here we present an approach for synthesizing biomedical knowledge based on a combination of word-embeddings and select cooccurrences. We evaluated our ability to recapitulate and retrospectively predict disease-gene associations from the Online Mendelian Inheritance in Man (OMIM) resource. Our metrics achieved an area under the curve (AUC) value of 0.981 at the recapitulation task for 2,400 disease-gene associations. At the most stringent cutoff, our metrics predicted 13.89% of these associations before their first cooccurrence in the literature, with a median time of 4 years between prediction and first cooccurrence. Finally, our literature metrics can be combined with human genetics data to retrospectively predict disease-gene associations, IL-6 and Giant Cell Arteritis provided as an example. We believe this framework can provide robust biomedical hypotheses at a much faster pace than current standard practices.

## Main Text

### Introduction

Biomedical knowledge, particularly in the form of literature text, has been growing at an increasing rate in recent years. For example, as of May 1^st^, 2019, PubMed contains 29,658,612 records, with 1,333,492 records from 2018 alone. While the explosion of biomedical knowledge has great potential in enabling new insights and discoveries, this same explosion of knowledge also makes it more difficult for scientists to read and process this wealth of information.

Databases such as ClinVar^1^ and BioGRID^2^ represent manually-curated efforts to summarize and store specific facets of biomedical knowledge (e.g. variation-phenotype associations and protein-protein interactions). Other tools parse the literature directly and utilize structured databases to extract important biomedical relationships in an automated manner. These tools include FACTA^3^ and PolySearch^4^. Finally, text-mining tools such as PubTator^5^ enable users to perform document triage, entity annotation, and relationship annotation in a PubMed-like interface.

In the last decade, substantial advances have been made in the field of natural language processing (NLP) that enable the rapid mining and extraction of relationships from biomedical literature. One such technique – word embeddings – leverages neural networks to capture the various semantic properties and relationships of words in a specified corpus of text. In 2013, Mikolov et al.^6^ introduced the word2vec model to produce distributed representations of words and phrases. In addition to word2vec, other word embedding models include GloVe^7^ and fastText^8,9^.

Pyysalo et al.^10^ were among the first to apply word2vec to biomedical literature, training their word2vec model on PubMed abstracts and PubMed Central full text papers. Minarro-Gimenez et al.^11^ subsequently trained a word2vec model on a corpus of biomedical literature that included PubMed, Merck Manuals, Medscape, and Wikipedia, and compared the model outputs to disease-drug relationships in the manually curated National Drug File – Reference Terminology (NDF-RT). In a follow-up work, Minarro-Gimenez et al.^12^ observed that the skip-gram model was more accurate than the continuous bag-of-words model, and that a window size of 10 and vector dimensionality of 300 produced the best results among all tested hyperparameter combinations. Bhasuran and Natarajan^13^ trained word2vec on a query-driven dataset of gene-disease associations from PubMed, in addition to gene-disease associations from four “gold standard corpora”. By combining word2vec output with other syntactic and semantic features, Bhasuran and Natarajan automatically extracted gene-disease associations from literature.

While the above models show the potential of word2vec for synthesizing biomedical knowledge, they also lack several important features: (1) the models lack a temporal dimension that reflects the movement of word vectors in semantic space over time as more knowledge is published; (2) the models do not address the existence of synonyms associated with biomedical entities (e.g. genes and diseases); (3) the standard metric for measuring the similarity of word vectors, the cosine distance, gives no indication if the association between the word vectors is statistically significant; (4) the models do not address the fact that knowledge is frequently siloed, so that associations found in one corpus of knowledge (e.g. scientific publications) are not necessarily found in other corpora (e.g. patents and SEC filings); (5) the models rely solely on word2vec and ignore traditional NLP techniques that could be used concurrently with word2vec; and (6) the models only incorporate unstructured knowledge (e.g. literature text) and do not incorporate structured knowledge such as human genetics datasets or RNA-Seq datasets.

Here we present a unified set of metrics to address the aforementioned shortcomings of current word2vec applications to biomedical literature. We report three distinct use cases for our metrics: (1) recapitulation of well-known disease-gene relationships using literature text; (2) retrospective prediction of well-known disease-gene relationships using literature text; and (3) retrospective prediction of disease targets using literature text and human genetics data from Ensembl^14,15^. For the first two use cases, we evaluated our approach against a positive set of disease-gene pairs from the Online Mendelian Inheritance in Man (OMIM) database^16-18^, and 15 similarly-sized negative sets of random disease-gene pairs. For the third use case, we evaluated our approach against the disease-target relationship between giant cell arteritis (GCA) and interleukin-6 (IL6) or the interleukin-6 receptor (IL6R). Our results indicate that our approach is able to recapitulate and retrospectively predict important biomedical associations using a combination of unstructured and structured data.

## Results

### Extracting biomedical associations from literature text

The approach we present is based on word embedding models trained on biomedical literature in various corpora and supplemented with named entity recognition, synonyms handling, and temporal slicing. Our approach uses two key metrics – the Local Score (*LS*) and Global Score (*GS*) – to quantify the strength of literature association between two concepts across various corpora. While *LS* measures how frequently two tokens are found within each other’s local context (the five tokens immediately preceding and following every occurrence of that token), *GS* measures how similar two tokens are when represented as word vectors in a high-dimensional semantic space. Each metric has its merits and drawbacks – for example, *LS* is useful for recapitulating well-known associations but is unable to identify novel associations. *GS*, on the other hand, is useful for identifying novel associations – particularly for concepts that have not cooccurred in the same document – but is unable to address synonyms for biomedical entities. *GS* is additionally unable to address polysemy (the coexistence of many possible meanings for a word or phrase) due to the context-independent nature of word2vec. In addition to incorporating unstructured knowledge such as literature text from various corpora, our approach can also incorporate structured knowledge such as human genetics, RNA-Seq, proteomics, and adverse event report datasets.

### Recapitulation of well-known disease-gene associations using literature text

We evaluated our ability to recapitulate well-known disease-gene associations using literature text against a positive set of 2,400 disease-gene pairs from OMIM and 15 negative sets. The disease-gene pairs in each negative set were generated by pairing the diseases from the positive set with random genes. We then calculated the percentage of disease-gene pairs in the positive and negative sets that satisfied either of the following conditions: (1) *LS* based on all knowledge produced up until today (*LS*_*today*_) was greater than or equal to a certain *LS* percentile cutoff or (2) *GS* based on all knowledge produced up until today (*GS*_*today*_) was greater than or equal to a certain *GS* percentile cutoff and had a nonzero *LS*_*today*_ (**Figure 1a**). Satisfying the former condition indicates that the disease and gene are frequently found within each other’s local context. Satisfying the latter condition, on the other hand, indicates that the word vectors corresponding to the disease and gene tokens are similar in a high-dimensional semantic space.

**Figure 1.**
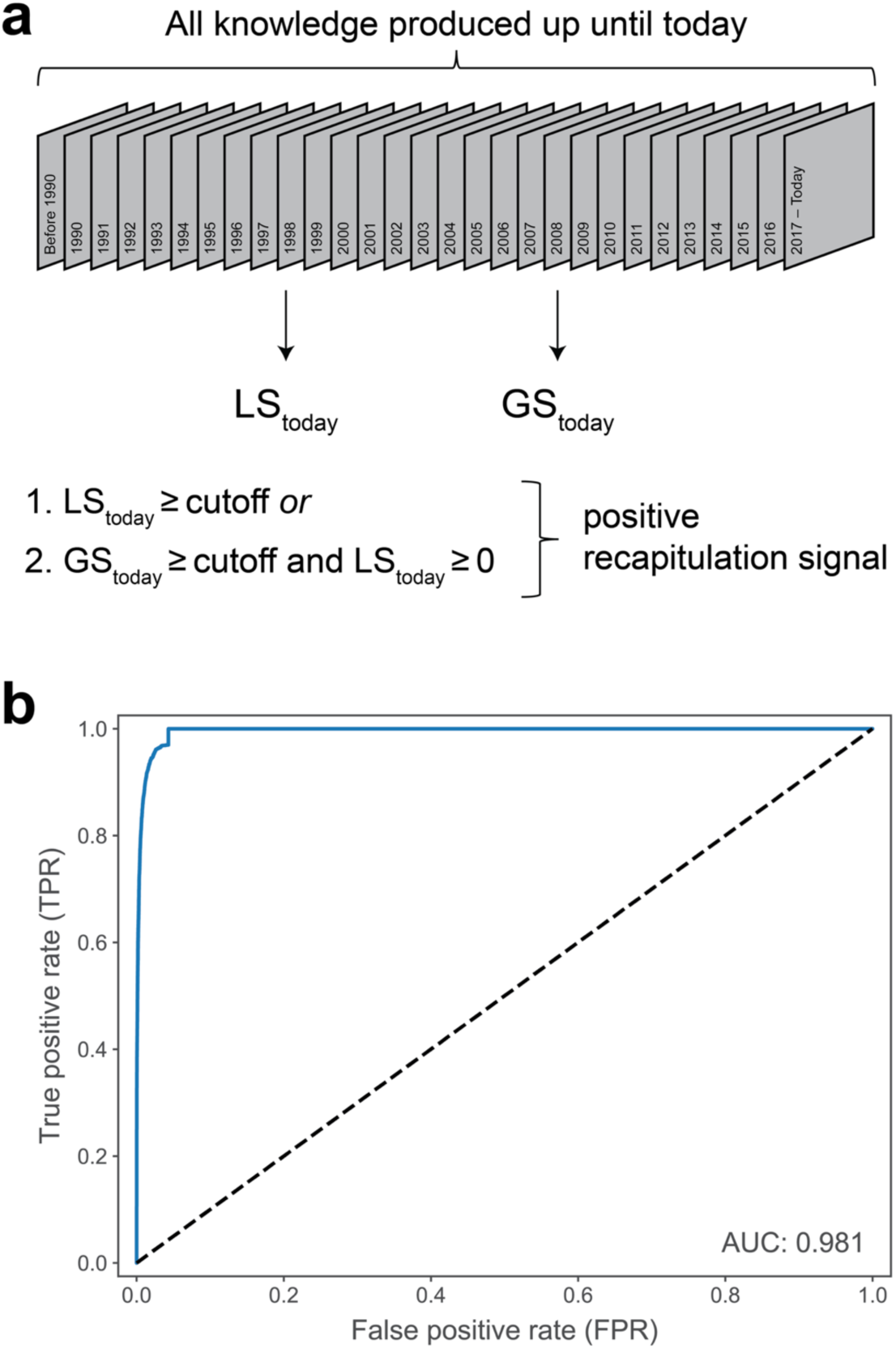
Recapitulation of well-known disease-gene associations using literature text. **(a)** *LS* and *GS* based on all knowledge produced up until today were calculated to evaluate our ability to recapitulate well-known disease-gene associations using literature text. **(b)** The receiver operating characteristic (ROC) curve, using the *LS* and *GS* percentile cutoff as the classification parameter. The true positive rate (TPR) is calculated from positive set of disease-gene pairs (n=2,249). The mean false positive rate (FPR) was calculated from the FPRs of all 15 negative sets of disease-gene pairs (n=2,397 to 2,400). The area under the curve (AUC) value was computed using the trapezoidal rule.

We calculated the true positive rate (TPR), mean false positive rate (FPR), and F_1_ score at various *LS* and *GS* percentile cutoffs between 0 and 100 (**Table 1 and Supplementary Table 2**). We plotted the receiver operating characteristic (ROC) curve using the *LS* and *GS* percentile cutoff as the classification parameter, and found that our approach achieved an area under the curve (AUC) value of 0.981 (**Figure 1b**). 96.98% of disease-gene pairs in the positive set have nonzero *LS*_*today*_or *GS*_*today*_, in contrast to 4.32% of disease-gene pairs in the negative sets (**Table 1**). It is important to note that we took the disease-gene associations from OMIM as ground truth. Because OMIM is extensively curated by scientific experts, the disease-gene associations reported by OMIM are most likely real. The absence of a disease-gene association in OMIM, however, does not exclude the possibility that the association is real. Nevertheless, our results indicate that our approach substantially recapitulates well-known disease-gene associations from literature text alone.

**Table 1.** Recapitulation of well-known disease-gene associations using literature. The true positive rate (TPR) is calculated from positive set of disease-gene pairs (n=2,249), and the number of predicted positive pairs is shown in parentheses. The mean and standard deviation of the false positive rate (FPR) are calculated from the FPRs of all 15 negative sets of disease-gene pairs (n=2,397 to 2,400).

| ***LS /GS***<br>**percentile**<br>**cutoff** | **TPR (n=2,249)** | **Mean FPR**<br>**(n=2,397 to**<br>**2,400)** | **Standard**<br>**deviation FPR**<br>**(n=2,397 to**<br>**2,400)** | **F<sub>1</sub> score** |
| --- | --- | --- | --- | --- |
| 99% | 73.01% (1,642) | 0.39% | 0.09% | 0.84 |
| 95% | 90.66% (2,039) | 1.27% | 0.21% | 0.94 |
| 90% | 94.53% (2,126) | 2.01% | 0.29% | 0.96 |
| 85% | 95.86% (2,156) | 2.59% | 0.32% | 0.97 |
| 80% | 96.44% (2,169) | 3.07% | 0.39% | 0.97 |
| 75% | 96.71% (2,175) | 3.38% | 0.39% | 0.97 |
| 50% | 96.98% (2,181) | 4.19% | 0.43% | 0.96 |
| 25% | 96.98% (2,181) | 4.32% | 0.43% | 0.96 |

**Table 2.** Retrospective prediction of well-known disease-gene associations using literature. The true positive rate (TPR) is calculated from positive set of disease-gene pairs (n=1,820), and the number of predicted positive pairs is shown in parentheses. The mean and standard deviation of the false positive rate (FPR) are calculated from the FPRs of all 15 negative sets of disease-gene pairs (n=2,397 to 2,400). The TPR and FPRs were computed using only the disease-gene pairs where the first year of cooccurrence happened after 1990, and the first year of cooccurrence happened after the first year of signal.

| **Signal consistency cutoff** | **TPR/FPR ratio** | **TPR (n=1,820)** | **Mean FPR (n=2,397 to 2,400)** | **Standard deviation FPR (n=2,397 to 2,400)** | **F<sub>1</sub> score** |
| --- | --- | --- | --- | --- | --- |
| 1% | 1.43 | 74.95% (1,364) | 52.29% | 1.02% | 0.61 |
| 10% | 1.94 | 71.37% (1,299) | 36.72% | 1.05% | 0.65 |
| 20% | 2.60 | 61.70% (1,123) | 23.74% | 0.89% | 0.64 |
| 30% | 3.29 | 50.00% (910) | 15.22% | 0.62% | 0.59 |
| 40% | 4.16 | 42.47% (773) | 10.20% | 0.57% | 0.54 |
| 50% | 5.11 | 35.49% (646) | 6.95% | 0.44% | 0.49 |
| 60% | 6.69 | 27.75% (505) | 4.15% | 0.30% | 0.42 |
| 70% | 8.74 | 22.31% (406) | 2.55% | 0.19% | 0.36 |
| 80% | 12.25 | 18.90% (344) | 1.54% | 0.16% | 0.31 |
| 90% | 18.98 | 14.51% (264) | 0.76% | 0.12% | 0.25 |
| 99% | 28.34 | 13.63% (248) | 0.48% | 0.11% | 0.24 |

### Retrospective prediction of well-known disease-gene associations using literature text

We next evaluated our ability to predict well-known disease-gene associations retrospectively using literature text against the same positive and negative sets of disease-gene pairs used above. For every disease-gene pair in the positive and negative sets, we computed a set of *GS* based on knowledge produced exclusively in each year between 1990 and 2017 (**Figure 2b**). We identified the first year where *GS* for a given disease-gene pair was nonzero, which we termed the first year of signal, and the first year where that disease and gene cooccurred in the PubMed corpus, which we termed the first year of cooccurrence (**Figure 2a**). Finally, we calculated the percentage of disease-gene pairs in the positive and negative sets where the signal consistency – the percentage of *GS* between the first year of signal and the first year of cooccurrence that were greater than or equal to 1.3 – was greater than or equal to a certain percentage cutoff (**Figure 2a**).

**Figure 2.**
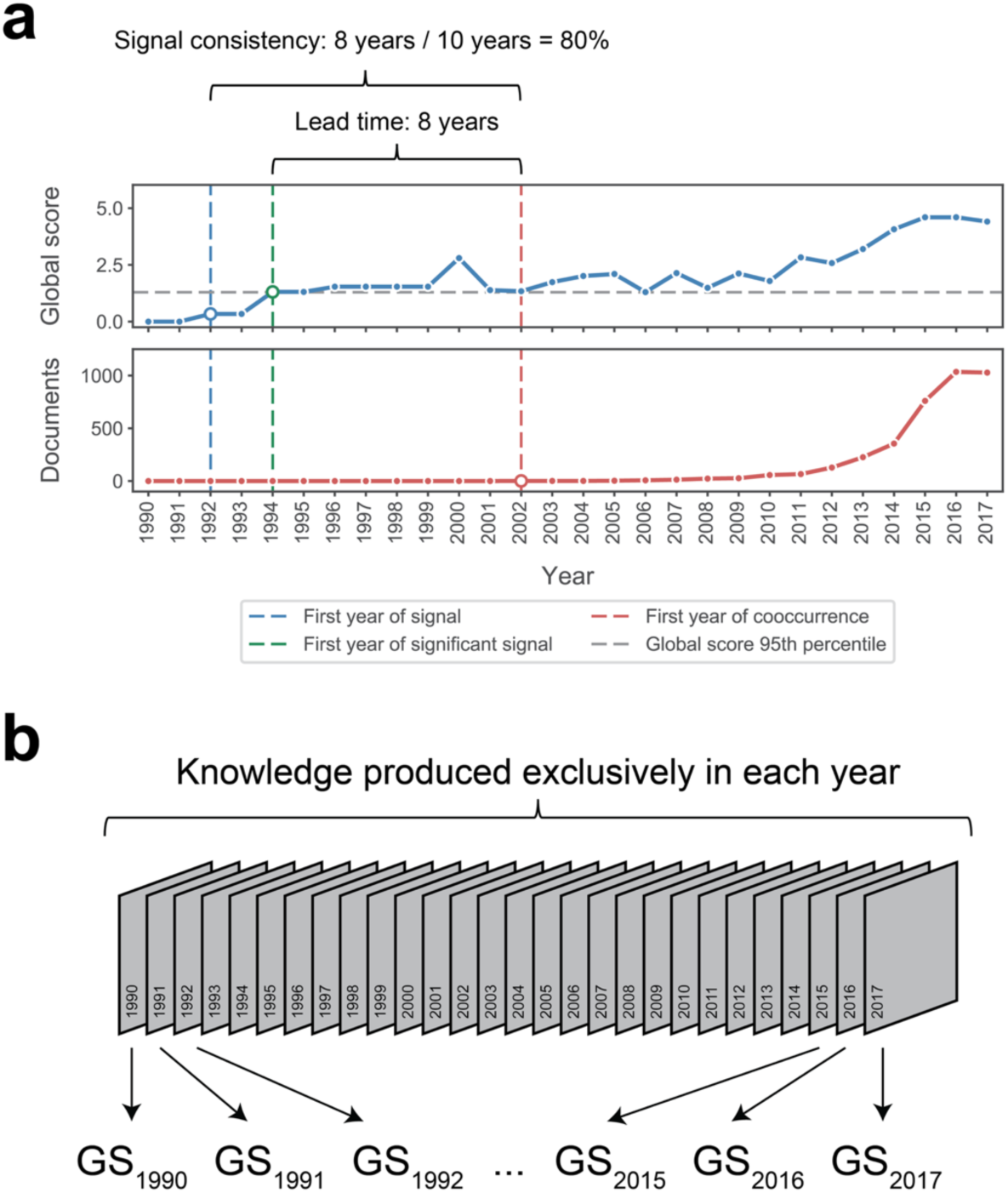
Retrospective prediction of well-known disease-gene associations using literature text. **(a)** Temporal evolution of *GS* and number of documents for the association between tumor-infiltrating lymphocytes and PD-L1 using the biomolecules collection as the control collection. The first year of signal is 1992, the first year of significant signal (*GS* ≥ 1.3) is 1994, and the first year of cooccurrence is 2002. Consequently, the lead time is 8 years and the signal consistency is 80%**. (b)** *GS* based on knowledge produced exclusively in each year between 1990 and 2017 were calculated to evaluate our ability to predict disease-gene associations retrospectively using literature text.

We calculated the TPR, mean FPR, TPR/FPR ratio, and F_1_ score at various signal consistency cutoffs between 0% and 100% (**Table 2**). As expected, we found that increasing the signal consistency cutoff decreased both the true positive rate (TPR) and the mean false positive rate (FPR) (**Figure 3a**). The mean FPR, however, decreased at a faster rate than the TPR when the signal consistency cutoff increased, as evidenced by the corresponding increase in the TPR/FPR ratio (**Figure 3b**). At our most stringent signal consistency cutoff (99%), our approach achieved a TPR of 13.63%, mean FPR of 0.48%, TPR/FPR ratio of 28.34, and F_1_ score of 0.24 (**Table 2**).

**Figure 3.**
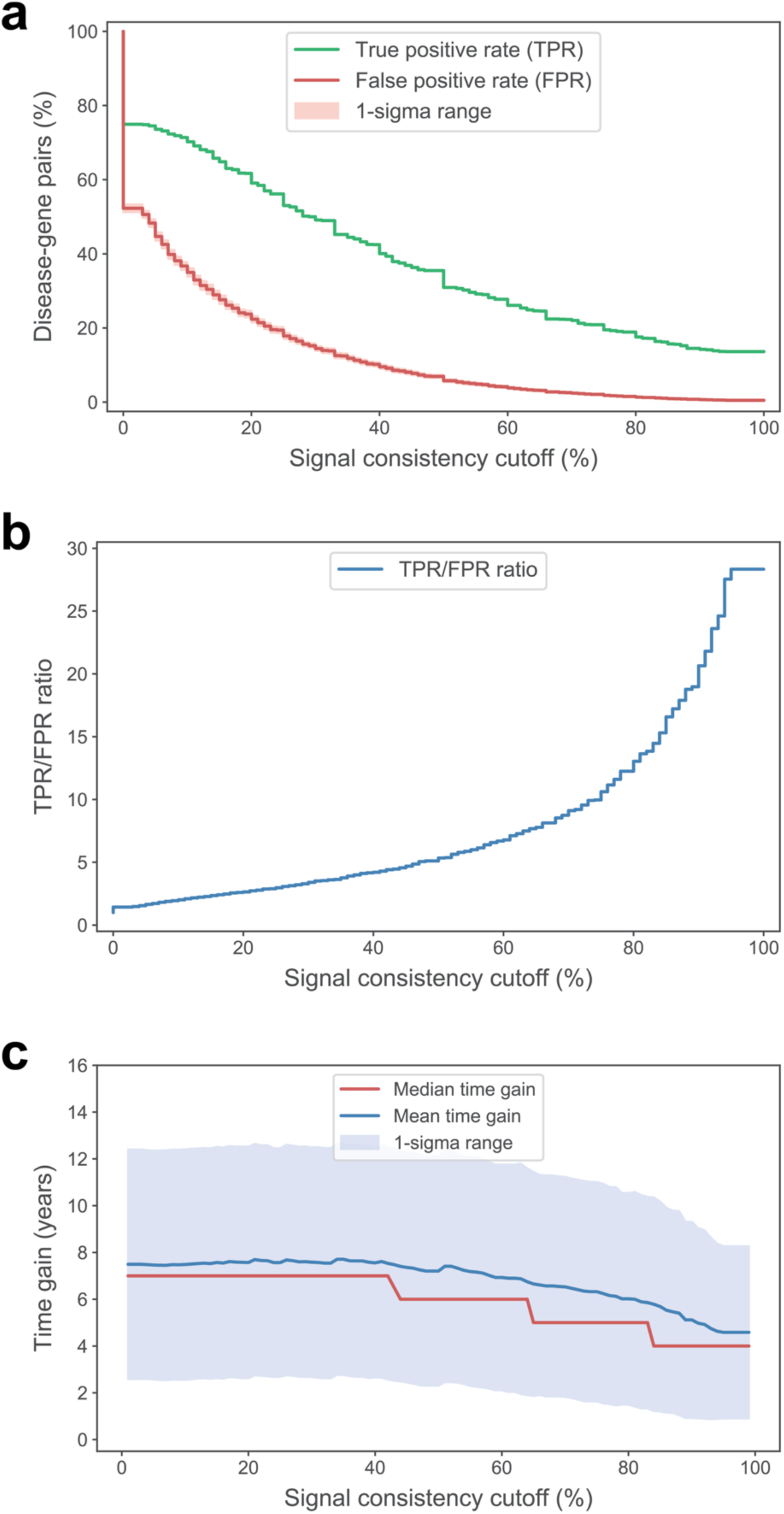
Retrospective prediction of well-known disease-gene associations using literature text. **(a)** Plot of the TPR (green) and mean FPR (red) as a function of the signal consistency cutoff. The TPR is calculated from positive set of diseasegene pairs (n=1,820). The mean and standard deviation of the FPR are calculated from the FPRs of all 15 negative sets of disease-gene pairs (n=2,397 to 2,400). The TPR and FPRs were computed using only the disease-gene pairs where the first year of cooccurrence happened after 1990, and the first year of cooccurrence happened after the first year of signal. **(b)** Plot of the TPR/FPR ratio as a function of the signal consistency cutoff. **(c)** Plot of the median (red), mean (blue), and 1-sigma range (light blue filled region) of the lead time as a function of the signal consistency cutoff. Lead times were calculated for the positive set of disease-gene pairs (n=1,820).

For each disease-gene pair in the positive set, we also identified the first year where *GS* for a given disease-gene pair was greater than 1.3, which we termed the first year of significant signal. We then calculated the lead time – the number of years between the first year of significant signal and the first year of cooccurrence (**Figure 2a**) – at various signal consistency cutoffs between 0% and 100% (**Table 3**). We found that the increasing the signal consistency cutoff decreased the median and mean lead times (**Figure 3c**). At our most stringent signal consistency cutoff (99%), our approach achieved a median lead time of 4.00 years and mean lead time of 4.58 years (**Table 3**).

**Table 3.**
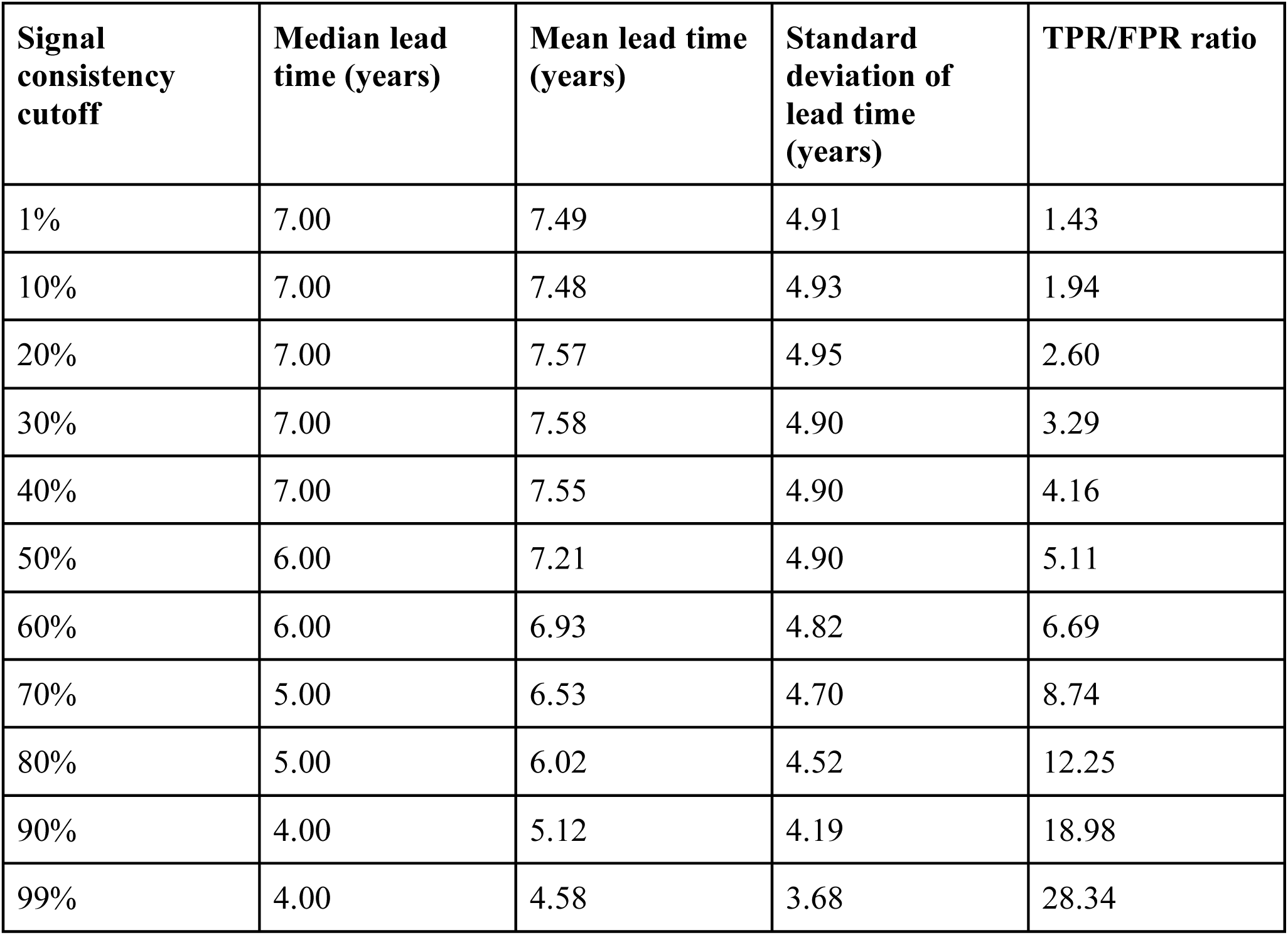
Retrospective prediction of well-known disease-gene associations using literature. The lead times were calculated from positive set of disease-gene pairs (n=1,820), and were computed using only the disease-gene pairs where the first year of cooccurrence happened after 1990, and the first year of cooccurrence happened after the first year of signal.

| **Signal consistency cutoff** | **Median lead time (years)** | **Mean lead time (years)** | **Standard deviation of lead time (years)** | **TPR/FPR ratio** |
| --- | --- | --- | --- | --- |
| 1% | 7.00 | 7.49 | 4.91 | 1.43 |
| 10% | 7.00 | 7.48 | 4.93 | 1.94 |
| 20% | 7.00 | 7.57 | 4.95 | 2.60 |
| 30% | 7.00 | 7.58 | 4.90 | 3.29 |
| 40% | 7.00 | 7.55 | 4.90 | 4.16 |
| 50% | 6.00 | 7.21 | 4.90 | 5.11 |
| 60% | 6.00 | 6.93 | 4.82 | 6.69 |
| 70% | 5.00 | 6.53 | 4.70 | 8.74 |
| 80% | 5.00 | 6.02 | 4.52 | 12.25 |
| 90% | 4.00 | 5.12 | 4.19 | 18.98 |
| 99% | 4.00 | 4.58 | 3.68 | 28.34 |

### Retrospective prediction of disease targets using literature text and human genetics data

Lastly, we evaluated our ability to predict disease targets retrospectively using literature text and human genetics data against the disease-target relationship between Giant Cell Arteritis (GCA) and IL6 or IL6R. We analyzed the temporal evolution of each target’s *LS* or *GS* rank to GCA relative to all genes, which represented our “literature-only approach” that did not incorporate any human genetics insights. We concurrently analyzed the temporal evolution of each target’s *LS* or *GS* rank to GCA relative to a set of genes with an indirect genetic association to GCA that changed every year, which represented our “literature plus genetics approach”. For the latter approach, we defined genes with an “indirect” genetic association to GCA as those with SNP evidence to phenotypes with *LS* ≥ 3.0 and *GS* ≥ 1.3 to GCA in that year.

We found that IL6’s *GS* rank from our literature-only approach first reached the 99th percentile in 2017, while IL6’s *GS* rank from our literature plus genetics approach first reached the 99th percentile in 2003 (**Figure 4a**). In addition, IL6’s *LS* rank from both approaches first reached the 99th percentile in 2006 (**Figure 4b**). Accordingly, the time of significance – defined as the number of years between the *GS* rank from our literature plus genetics approach first reaching the 99th percentile and the *LS* rank from our literature-only approach first reaching the 99th percentile – was 3 years(**Table 4**).

**Table 4.** Temporal evolution of IL6’s *LS* and *GS* ranks to GCA using the literature-only and literature plus genetics approaches.

| **Year** | ***LS* only** | ***LS* and genetics** | ***GS* only** | ***GS* and genetics** |
| --- | --- | --- | --- | --- |
| 1990 | 13,839 | 13,839 | 13,839 | 13,839 |
| 1991 | 13,839 | 13,839 | 1,668 | 13,839 |
| 1992 | 13,839 | 13,839 | 13,839 | 13,839 |
| 1993 | 13,839 | 13,839 | 442 | 13,839 |
| 1994 | 13,839 | 13,839 | 2,489 | 13,839 |
| 1995 | 13,839 | 13,839 | 13,839 | 13,839 |
| 1996 | 13,839 | 13,839 | 2,127 | 13,839 |
| 1997 | 13,839 | 13,839 | 1,896 | 13,839 |
| 1998 | 13,839 | 13,839 | 923 | 13,839 |
| 1999 | 13,839 | 13,839 | 1,370 | 13,839 |
| 2000 | 13,839 | 13,839 | 2,587 | 13,839 |
| 2001 | 13,839 | 13,839 | 317 | 13,839 |
| 2002 | 13,839 | 13,839 | 1,222 | 13,839 |
| 2003 | 13,839 | 360 | 1,888 | 64 |
| 2004 | 13,839 | 506 | 1.472 | 74 |
| 2005 | 13,839 | 721 | 209 | 24 |
| 2006 | 14 | 4 | 857 | 115 |
| 2007 | 8 | 2 | 1,166 | 112 |
| 2008 | 9 | 2 | 555 | 58 |
| 2009 | 12 | 2 | 1,538 | 187 |
| 2010 | 7 | 1 | 292 | 68 |
| 2011 | 10 | 3 | 330 | 68 |
| 2012 | 6 | 3 | 257 | 49 |
| 2013 | 4 | 2 | 259 | 46 |
| 2014 | 4 | 2 | 452 | 87 |
| 2015 | 4 | 2 | 637 | 123 |
| 2016 | 4 | 2 | 208 | 43 |
| 2017 | 4 | 2 | 63 | 20 |
| Today | 4 | 2 | 100 | 29 |

**Figure 4.**
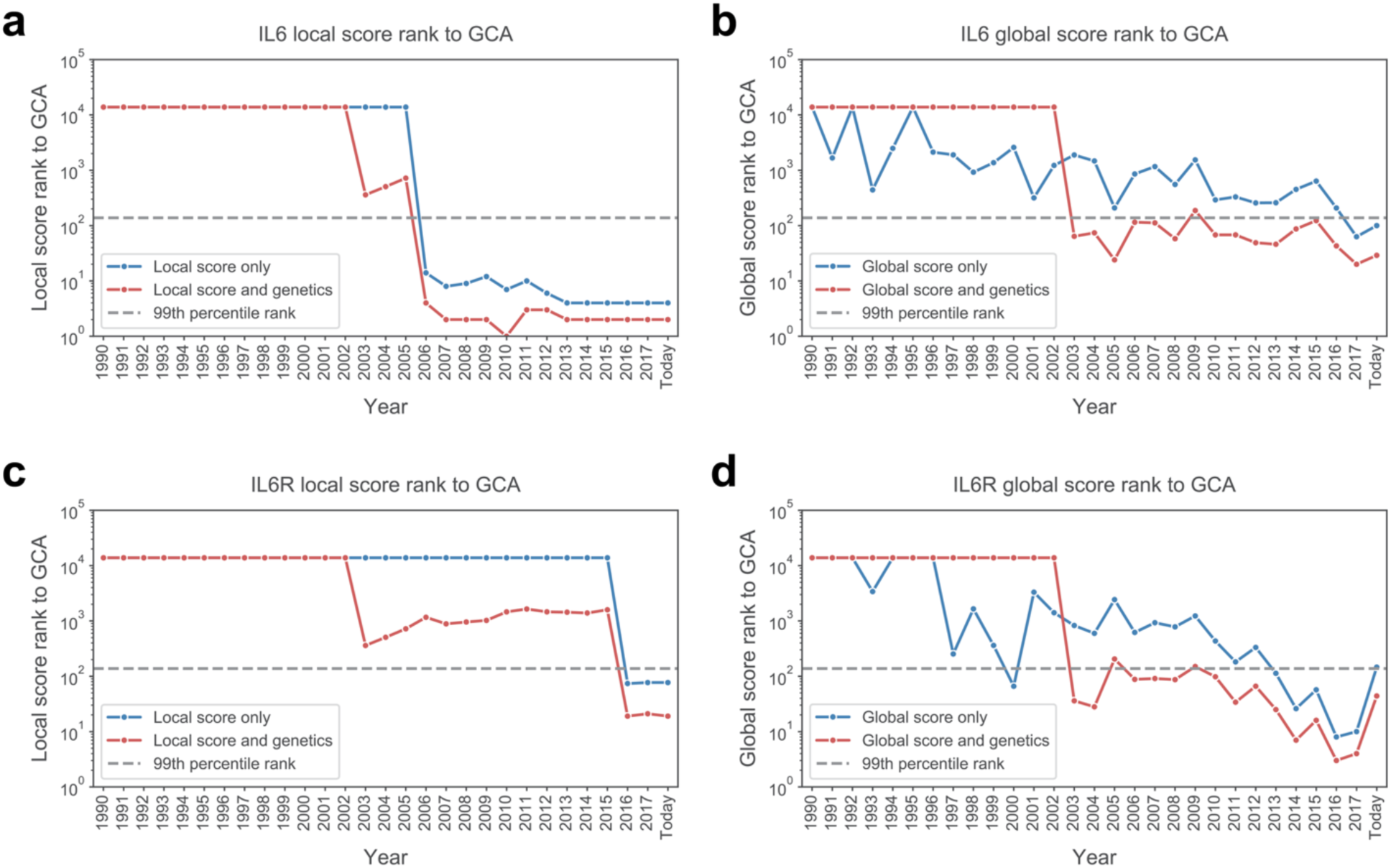
Temporal evolution of IL6 and IL6R’s *LS* and *GS* ranks to GCA using the literature text and human genetics data. **(a)** Plot of the temporal evolution of IL6’s *LS* rank to GCA using the literature-only (blue) and literature plus genetics (red) approaches. **(b)** Plot of the temporal evolution of IL6’s *GS* rank to GCA using the literature-only (blue) and literature plus genetics (red) approaches. **(c)** Plot of the temporal evolution of IL6R’s *LS* rank to GCA using the literature-only (blue) and literature plus genetics (red) approaches. **(d)** Plot of the temporal evolution of IL6R’s *GS* rank to GCA using the literature-only (blue) and literature plus genetics (red) approaches.

We found that while IL6R’s *GS* rank from our literature-only approach first reached the 99th percentile in 2000, the rank did not consistently exceed the 99th percentile until 2013 onwards (**Figure 4c**). IL6R’s *GS* rank from our literature plus genetics approach, on the other hand, first reached the 99th percentile in 2003 (**Figure 4c**). In addition, IL6R’s *LS* rank from both approaches first reached the 99th percentile in 2016 (**Figure 4d**). Accordingly, the time of significance between the *GS* rank from the literature plus genetics approach reaching the 99th percentile and the *LS* rank from the literature-only approach reaching the 99th percentile was 13 years (**Table 5**).

**Table 5.** Temporal evolution of IL6R’s *LS* and *GS* ranks to GCA using the literature-only and literature plus genetics approaches.

| **Year** | ***LS* only** | ***LS* and genetics** | ***GS* only** | ***GS* and genetics** |
| --- | --- | --- | --- | --- |
| 1990 | 13,839 | 13,839 | 13,839 | 13,839 |
| 1991 | 13,839 | 13,839 | 13,839 | 13,839 |
| 1992 | 13,839 | 13,839 | 13,839 | 13,839 |
| 1993 | 13,839 | 13,839 | 3,396 | 13,839 |
| 1994 | 13,839 | 13,839 | 13,839 | 13,839 |
| 1995 | 13,839 | 13,839 | 13,839 | 13,839 |
| 1996 | 13,839 | 13,839 | 13,839 | 13,839 |
| 1997 | 13,839 | 13,839 | 253 | 13,839 |
| 1998 | 13,839 | 13,839 | 1,650 | 13,839 |
| 1999 | 13,839 | 13,839 | 361 | 13,839 |
| 2000 | 13,839 | 13,839 | 66 | 13,839 |
| 2001 | 13,839 | 13,839 | 3,305 | 13,839 |
| 2002 | 13,839 | 13,839 | 1,407 | 13,839 |
| 2003 | 13,839 | 360 | 827 | 36 |
| 2004 | 13,839 | 506 | 597 | 28 |
| 2005 | 13,839 | 721 | 2,426 | 205 |
| 2006 | 13,839 | 1,163 | 620 | 88 |
| 2007 | 13,839 | 886 | 928 | 91 |
| 2008 | 13,839 | 956 | 782 | 87 |
| 2009 | 13,839 | 1,022 | 1,230 | 150 |
| 2010 | 13,839 | 1,457 | 434 | 98 |
| 2011 | 13,839 | 1,641 | 182 | 34 |
| 2012 | 13,839 | 1,459 | 332 | 66 |
| 2013 | 13,839 | 1,437 | 113 | 25 |
| 2014 | 13,839 | 1,391 | 26 | 7 |
| 2015 | 13,839 | 1,588 | 57 | 16 |
| 2016 | 74 | 19 | 8 | 3 |
| 2017 | 77 | 21 | 10 | 4 |
| Today | 77 | 19 | 146 | 44 |

## Discussion

The unified approach we present incorporates both unstructured and structured data to synthesize large amounts of biomedical knowledge. The *LS* and *GS* metrics based on all knowledge produced up until today substantially recapitulate disease-gene relationships from OMIM and complement each other in various aspects. In addition, the *GS* metric based on knowledge produced exclusively in each year between 1990 and 2017 can be strongly predictive of disease-gene relationships, even before the first cooccurrence of the disease and gene in the PubMed corpus, a power we believe could extend to scores obtained today. These two results demonstrate our approach’s utility to users who are interested in (1) well-known biomedical associations in unfamiliar subject areas, or (2) novel biomedical associations in familiar subject areas. Importantly, word embedding models such as word2vec can identify semantic associations between two concepts that cooccur only a few times or never cooccur at all in a given corpus of text. The cosine distance metric most frequently used in word embedding models, however, is not indicative of the statistical significance of any given semantic association. Our approach addresses this limitation of the cosine distance by introducing a control collection – a group of tokens belonging to a shared entity type (e.g. diseases) – and computing *GS* to indicate the statistical significance of a semantic association between two concepts. *LS*, on the other hand, is a modified variant of the pointwise mutual information metric that only considers tokens that are in each other’s local context – the five tokens immediately preceding and following every occurrence of that token.

As noted previously, context-independent word embedding models such as word2vec are unable to address polysemy. For example, word2vec is unable to discern that the biomedical term “EGFR” could refer to either the gene known as “epidermal growth factor receptor” or the clinical measurement known as “estimated glomerular filtration rate”. Newer context-sensitive models such as ELMo^19^ and transformed-based models such as BERT^20^ and GPT-2^21^ are able to address polysemy. We are actively working on the implementation of these models for future versions of our approach, and accordingly updating the *GS* metric to incorporate polysemy.

While evaluating our ability to predict well-known disease-gene associations retrospectively using literature text, we observed that both the TPR/FPR ratio increases and median/mean lead time decreases with increasing signal consistency cutoffs. The TPR/FPR ratio reflects the enrichment of actual disease-gene associations compared to random diseasegene associations. The lead time, on the other hand, reflects how many years in advance our approach detected a significant *GS* for an actual disease-gene association before that disease and gene cooccurred in the same document. Concurrently, the TPR – which reflects the fraction of actual disease-gene associations that are correctly identified by our approach – decreases with increasing signal consistency cutoffs. A user interested in using our approach for a hypothesis generation project (e.g. target identification) consequently must balance two opposing trends when picking an appropriate signal consistency cutoff. Increasing the enrichment of actual disease-gene associations compared to random disease-gene associations comes at the expense of not only incorrectly throwing out a greater proportion of actual disease-gene associations, but also decreasing the median/mean lead time. An appropriate signal consistency cutoff is therefore determined by a combination of the user’s end goals and risk tolerance.

Our signal consistency and lead time calculations both depended on the first year the disease and gene cooccurred in the PubMed corpus (“first year of cooccurrence”), rather than the first year the disease and gene were definitively linked in a publication (“first year of discovery”). Since the first year of cooccurrence is usually earlier than the first year of discovery, our decision to use the former to evaluate our approach’s performance at the retrospective prediction task represents a more conservative approach. From the user’s perspective, the first year of discovery might be more relevant than the first year of cooccurrence when calculating the signal consistency and lead time metrics, since a disease-gene association most likely would become actionable only after a definitive link has been published. Nevertheless, our results using the first year of cooccurrence demonstrate that our approach can retrospectively predict disease-gene relationships well before the first year of cooccurrence and the first year of discovery.

We generated our positive set of 2,400 disease-gene pairs by taking disease-gene associations from OMIM as ground truth for two main reasons. First, OMIM is comprehensive and extensively curated by scientific experts. Second, the diseases represented in OMIM are mainly monogenic and/or Mendelian. We consequently generated negative sets of diseases paired with random genes in order to evaluate our ability to discriminate between signal – actual disease-gene associations – and noise – random disease-gene associations. Because most genes are related to one another via a complex network of biological pathways, it is difficult to define a “true negative” in the context of disease-gene associations. For example, while mutations in a single gene might be responsible for a disease (e.g. *HEXA* mutations in Tay-Sachs disease), the disease usually manifests itself via disruptions in the numerous biological pathways that the mutated gene is involved in (e.g. ganglioside metabolism). Nevertheless, we believe that pairing a set of mainly monogenic and/or Mendelian diseases with random genes represents the best way of generating “true negatives” without introducing additional complexities.

We additionally demonstrated that our approach can combine literature text with knowledge from structured datasets such as human genetics data. While insights generated by our approach based solely on literature text are useful for hypothesis generation projects, insights generated by our approach based on literature text and other structured data are useful for hypothesis prioritization projects. In the case of the disease-target relationship between GCA and IL6 or IL6R, introducing human genetics data winnowed out genes that were genetically unrelated to GCA, enabling both IL6 and IL6R to emerge as relevant hits. In addition, we introduced the concept of “indirect” genetic relationships based on semantically related phenotypes to GCA, which allowed us to gain significant lead time in the association. Introducing additional knowledge siloes, whether structured or unstructured, enable the user to triangulate concordant signals that exist across all siloes and accordingly prioritize hypotheses that have strongest concordance.

In summary, we have described an approach for synthesizing biomedical knowledge and report three distinct use cases. Importantly we have incorporated a temporal dimension that we have used to demonstrate that our metrics can retrospectively predict well-known disease-gene relationships. Moreover, we can additionally incorporate structured data such as human genetic associations (i.e. GWAS) to triangulate insights from literature text, which we have used to demonstrate that our approach can predict the disease-target relationship between GCA and IL6 or IL6R before it became known in the literature. In the future we intend to adapt our metrics to include context-sensitive word-embeddings, and to apply our methods to other biomedical corpora outside of the main scientific literature, most importantly Electronic Health Records (EHR), and to triangulate insights with larger amounts of structured data (e.g. next-generation sequencing).

## Methods

### Data ingestion and knowledge extraction pipeline

Data was ingested via web crawlers or downloaded directly from sources. The text was extracted from the ingested data, and phrase generation was performed to extract multi-word tokens (e.g. “non-small-cell lung cancer”) that occur frequently enough to warrant their own embedding. A “token” refers to each individual concept (word or phrase) that is part of our knowledge graph. The phrase generation approach was based on a neural network model for part-of-speech tagging^22,23^. Embeddings were generated from the extracted text using word2vec (https://github.com/tmikolov/word2vec) and the skip-gram model with negative sampling and the following hyperparameters: negative sample size of 5, sub-sampling of 1e-5, minimum count of 7, learning rate of 0.025, vector dimensionality of 300, and window size of 5. Separate embeddings were generated for the various corpora consumed by our approach, and for various time slices (every year from 1990 to 2017). Each embedding was generated from randomly initialized vectors, such that embeddings from one time slice (e.g. 2010) were independent of all other time slices (e.g. 1990 to 2009, 2011 to 2017).

### Sources consumed

Our knowledge graph supports embeddings for various corpora. **Supplementary Table 1** details the different corpora we have currently consumed, including the number of documents, words, and vectors within each corpus.

### Key metrics: Local Score (*LS*)

The Local Score (*LS*) measures how frequently two tokens are found within each other’s local context in a corpus, normalized by the occurrences of those tokens in that corpus. We define the local context of a token as the five tokens immediately preceding and following every occurrence of that token. We additionally define the adjacency adj_AB_ between tokens A and B as the number of times token A is found token B’s local context, or vice-versa. We calculate the pointwise mutual information pmi_AB_ between tokens A and B as the following:

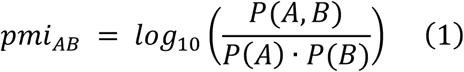

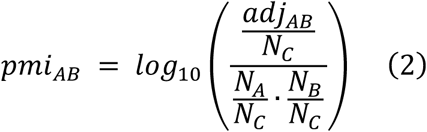

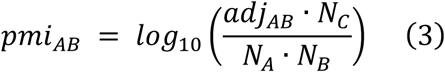

Where *P*(*A, B*) is the probability of observing tokens A and B in each other’s local context, *P*(*A*) is the probability of observing token A, *P*(*B*) is the probability of observing token B, N_A_ is the occurrences of token A, N_B_ is the occurrences of token B, and N_C_ is the summed occurrences of all tokens in the corpus of interest. We then calculate LS_AB_, the *LS* between tokens A and B as the following:

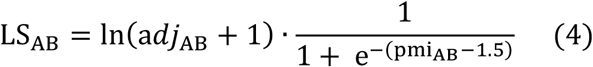

One disadvantage we observed of using pmi_AB_ on its own is that rarely occurring tokens tend to dominate the measure. For example, let token A with N_A_ = 1 have its only occurrence be in the local context of token B with N_B_ = 1.0 × 10^6^. If N_C_ = 1.0 × 10^11^ then pmi_AB_= 5.0. Our formula for LS_AB_ passes pmi_AB_ through a shifted sigmoid function (with an arbitrary midpoint of 1.5 chosen based on performance) to dampen large values of pmi_AB_. The Ln(*adj*_AB_ + 1) term additionally down-weights rarely occurring tokens relative to more commonly occurring tokens. Therefore, LS_AB_ for tokens A and B in our hypothetical scenario would be 0.67. We have empirically observed that *LS* is approximately distributed according to an exponential distribution with rate parameter λ = 1.0 (**Supplementary Figure 1**), and the percentile rank conversion table for *LS* in our approach is shown in **Supplementary Table 2**. We define significant associations as those having a *LS* ≥ 3.0, representing the top 5% of associations in this exponential distribution.

### Key metrics: Composite Local Score (Composite *LS*)

While *LS* measures how frequently two tokens are found within each other’s local context, the Composite Local Score (Composite *LS*) measures how frequently two groups of tokens are found within each other’s local context. Consequently, the composite *LS* is particularly useful for handling token synonyms. We calculate the composite pointwise mutual information 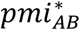 as the following:

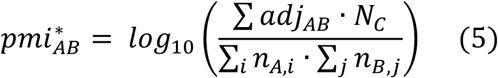

Where Σ_*i*_ *n*_*A,i*_ is the summed occurrences of token A and its synonyms, and Σ_*j*_ *n*_*B,j*_ is the summed occurrences of token B and its synonyms, and Σ *adj*_*AB*_ is the summed adjacency between tokens A and B and their respective synonyms. We then calculate 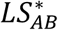, the Composite *LS* between tokens A and B and their respective synonyms as the following:

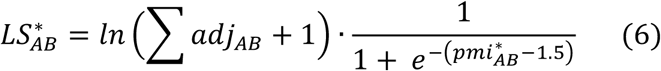

### Key metrics: Global Score (*GS*)

The Global Score (*GS*) measures the similarity between the word vectors – or numerical representations in a high-dimensional semantic space – corresponding to two tokens, normalized by a control token collection – or group of tokens belonging to a shared category (e.g. diseases). We calculate the cosine distance *D*_*AB*_ between tokens A and B, which are members of token collections *C*_*A*_ and *C*_*B*_ respectively, as the following:

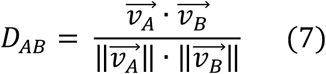

Where 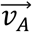 is the word vector corresponding to token A and 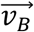 is the word vector corresponding to token B. Because the cosine distance *D*_*AB*_ in isolation does not indicate if the association between tokens A and B is statistically significant, we must consider how the cosine distances between token A and all tokens in token collection *C*_*B*_ are distributed, or how the cosine distances between token B and all tokens in token collection *C*_*A*_ are distributed. We calculate 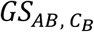, the *GS* between tokens A and B with control token collection *C*_*B*_ as the following:

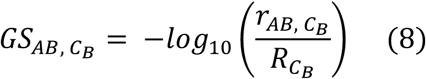

Where 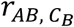 is the rank of *D*_*AB*_ relative to all cosine distances between token A and all tokens in *C*_*B*_, and 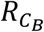 is the size of *C*_*B*_. Because *GS* depends on the control token collection, it is important to note that:

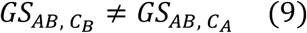

Unless *C*_*A*_ and *C*_*B*_ are the same token collection. *D*_*AB*_ is approximately distributed according to a normal distribution (**Supplementary Figure 2**), and the percentile conversion table for *GS* in our approach is shown in **Supplementary Table 2**. We define significant associations as those having a *GS* ≥ 1.3, representing the top 5% of associations in the normal distribution for all cosine distances between 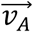 and all tokens in *C*_*B*_.

### Extracting disease-gene pairs from the Online Mendelian Inheritance in Man (OMIM) resource

We used the Online Mendelian Inheritance in Man (OMIM) Gene Map to extract a list of validated disease-gene associations. The OMIM Gene Map describes associated genes and phenotypes for 16,839 unique cytogenetic locations described in OMIM and previously published in cited references. For every gene represented in the OMIM Gene Map, we extracted all its associated phenotypes. We then removed any phenotypes that were flagged by OMIM as a non-disease (e.g. “lean body mass”) or susceptibility phenotype (e.g. “resistance to malaria”), in addition to phenotypes deemed by OMIM as having a provisional relationship with that gene. Consequently, we extracted a total of 4,996 disease-gene pairs – which we termed “OMIM disease-gene pairs” – comprising 4,653 unique diseases and 3,608 unique genes.

### Mapping disease and gene names to relevant tokens

For every disease represented in the OMIM disease-gene pairs, we mapped the full disease name provided by OMIM to the relevant preferred name token for that disease. Mapping to the relevant preferred name token was necessary since tokens serve as the input to queries in our approach, and the preferred name token was required for extracting synonym tokens. When we were unable to map the full disease name, we attempted to map part of the full disease name to the relevant preferred name token (e.g. we mapped “rhabdomyosarcoma, embryonal, 1” to “embryonal rhabdomyosarcoma”). After identifying the preferred name token, we extracted all synonym tokens that were associated to that disease. We then removed any synonym tokens with a fewer than 1,000 occurrences in the core corpus, since we have observed that *GS* for tokens with fewer than 1,000 occurrences are insufficiently robust. We additionally removed synonym tokens that were shared with other diseases or genes, or were otherwise ambiguous. In total, we identified the disease preferred name token for 3,487 disease-gene pairs comprising 3,187 unique diseases and 2,618 unique genes.

For every gene represented in the OMIM disease-gene pairs, we similarly identified the preferred name token for that gene and all synonym tokens that were associated to that gene, including the HGNC symbol. We removed all synonym tokens with fewer than 1,000 occurrences in the core corpus, and all ambiguous synonym tokens. In total, we identified the gene preferred name token for 4,986 disease-gene pairs, comprising 4,643 unique diseases and 3,601 unique genes.

### Identifying the first year of cooccurrence for disease-gene pairs

For every OMIM disease-gene pair, we additionally identified the first year where the disease (or any of its synonyms) cooccurred with the gene (or any of its synonyms) in the PubMed corpus, which we termed the “first year of cooccurrence”. We identified the first year of cooccurrence for 3,395 disease-gene pairs comprising 3,097 unique diseases and 2,541 unique genes.

### Defining a positive set of disease-gene pairs

We defined a positive set of disease-gene pairs by taking all OMIM disease-gene pairs where the disease and gene preferred name tokens both had greater than 1,000 occurrences in the core corpus. In total, there were 2,400 disease-gene pairs that met these criteria, which comprised 2,138 unique diseases, 683 unique disease preferred name tokens, and 1,799 unique genes. Since multiple diseases could map onto the same disease preferred name token (e.g. we mapped both “spinal muscular atrophy, type I” and “spinal muscular atrophy, type II” to “spinal muscular atrophy”), there were 2,249 unique pairs of disease and gene preferred name tokens.

### Defining a negative set of disease-gene pairs

We defined a negative set of disease-gene pairs by fixing the diseases in our positive set and pairing them with genes that were randomly drawn from a set of 11,544 genes whose gene HGNC tokens had greater than 1,000 occurrences in the core corpus. We generated 15 negative sets comprising 2,400 disease-gene pairs.

### Computing composite *LS* for disease-gene pairs

We computed the composite *LS* for each disease-gene pair using the disease preferred name token and the gene synonym tokens based on all knowledge produced up until today in the core corpus.

### Computing *GS* for disease-gene pairs

We computed *GS* for each disease-gene pair for multiple time slices using the disease preferred name token and gene synonym tokens. First, we computed *GS* based on all knowledge produced up until today in the core corpus. Second, we computed *GS* based on knowledge produced exclusively in each year between 1990 (the earliest year for which *GS* is available) and 2017 (the most recent year for which *GS* is available).

Because the *GS* calculation depends on the control collection, we computed each *GS* using the diseases and genes collections as controls and took the maximum of the two values. In addition, when there were multiple synonym tokens for a gene, we computed *GS* for each disease-gene synonym pair and took the maximum value.

The *GS* for a disease-gene pair is not available for years in which either token does not exceed a minimum number of occurrences. In these cases, we used the *GS* from the most recent year for which *GS* was available.

### Evaluating recapitulation of well-known disease-gene associations using literature text

To evaluate our ability to recapitulate well-known disease-gene associations using literature text, we considered two metrics for each disease-gene pair – (1) *GS*_*today*_, the *GS* based on all knowledge produced up until today and (2) *LS*_*today*_, the composite *LS* based on all knowledge produced up until today. We deemed a disease-gene pair to have a positive signal if it satisfied one of the two following conditions: (1) *LS*_*today*_was greater than or equal to a certain percentile cutoff (e.g. 3.0 for the 95th percentile) or (2) *GS*_*today*_was greater than or equal to a certain percentile cutoff (e.g. 1.3 for the 95th percentile) and had a nonzero *LS*_*today*_.

### Evaluating retrospective prediction of well-known disease-gene associations using literature text

To evaluate our ability to predict well-known disease-gene associations retrospectively using literature text, we first computed the signal consistency, defined as the percentage of *GS* between the first year of signal and the first year of cooccurrence that were greater than or equal to 1.3. For disease-gene pairs in the negative set that did not have a first year of cooccurrence, the signal consistency was defined as the percentage of *GS* between the first year of signal and 2017 (the latest year for which *GS* is available) that were greater than or equal to 1.3. We deemed a disease-gene pair to have a positive signal if the signal consistency a certain percentage cutoff. Because the earliest year for which *GS* is available was 1990, we computed the signal consistency only for disease-gene pairs where the first year of cooccurrence happened after 1990, and the first year of cooccurrence happened after the first year of signal.

We additionally computed the lead time – defined as the number of years between the first year *GS* was greater than or equal to 1.3 and the first year of cooccurrence – for the diseasegene pairs in our positive set. We similarly computed the lead time only for disease-gene pairs where the first year of cooccurrence happened after 1990, and the first year of cooccurrence happened after the first year of signal.

### Evaluating retrospective prediction of disease targets using literature text and human genetics data

To evaluate our ability to predict disease targets retrospectively using literature text and human genetics data, we computed the *LS* between GCA and a set of 13,839 genes that had at least one synonym token with greater than 1,000 occurrences in the core corpus based on knowledge produced up until every year between 1990 and 2017. We additionally computed the *GS* between GCA and the same set of 13,839 genes based on knowledge produced exclusively in each year between 1990 and 2017. We performed similar *LS* and *GS* computations between GCA and a set of 20,893 human disease phenotypes.

We then computed IL6’s or IL6R’s *LS* or *GS* rank to GCA relative to two different gene sets. First, we computed the *LS* or *GS* rank relative to all 13,839 genes, which represented a “literature-only approach”. Second, we computed the *LS* or *GS* rank relative to genes that had an indirect genetic association to GCA in each year before 1990 and 2017, which represented a “literature plus genetics approach”. We defined genes with an indirect genetic association to GCA as genes with human SNP evidence (according to Ensembl) to phenotypes with *LS* ≥ 3.0 and *GS* ≥ 1.3 to GCA.

## Acknowledgments

The authors thank Karthik Murugadoss, Sankar Ardhanari, Venky Soundararajan, Prashanth Ellina, and Christopher Sarkisian for important and rewarding discussions.

## Author Contributions Statement

E.G.R., A.L.M. and M.A conceived the study and analyses. A.R., A.P., and M.A derived and implemented the *LS* and *GS* metrics. J.P. acquired the data and executed the study. J.P., A.L.M., and E.G.R. analyzed and interpreted the data. J.P. and E.G.R. wrote the manuscript, with input from all authors.

## Supplementary Information

**Supplementary Figure 1.**
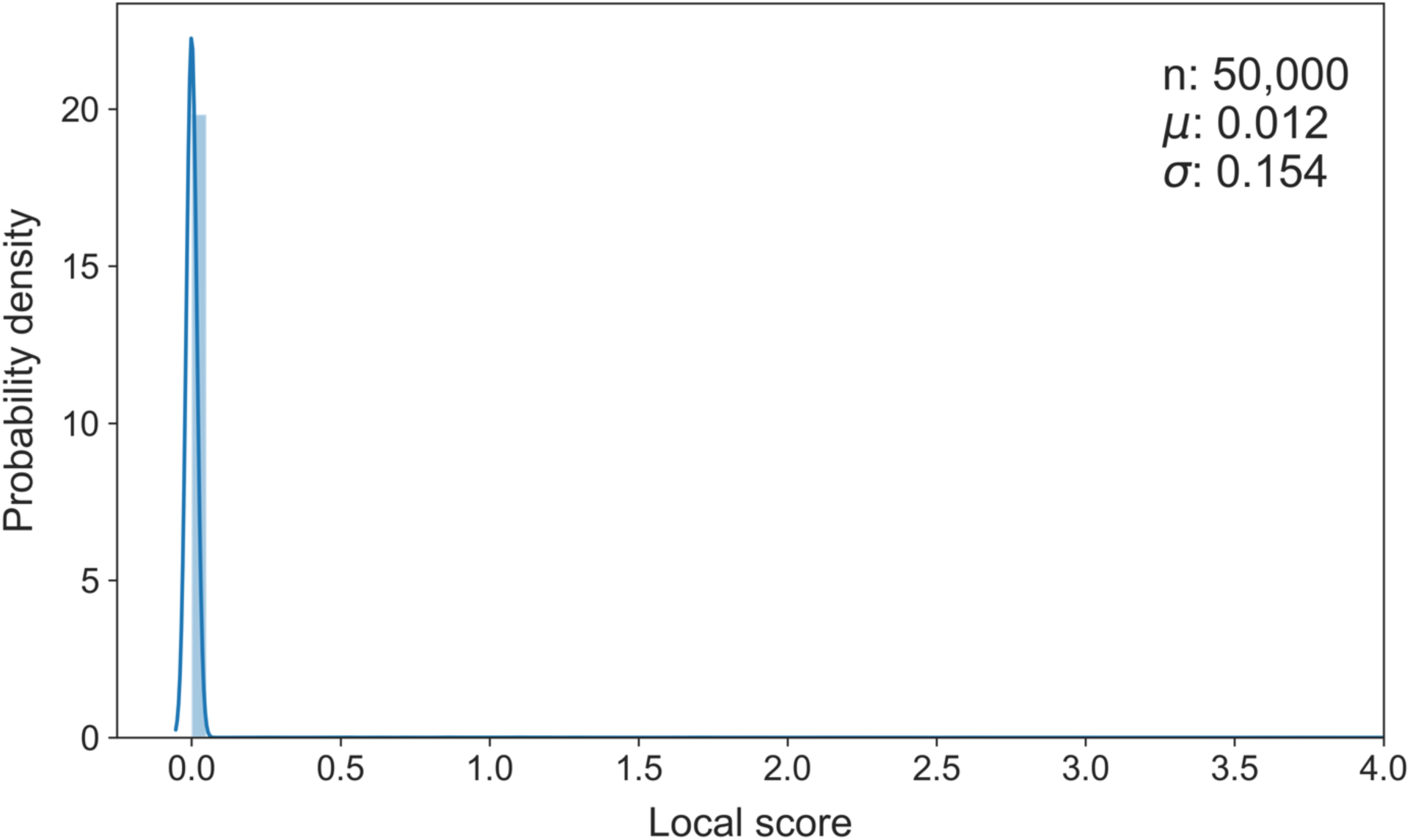
Distribution of local scores. Local scores were calculated from 50,000 pairs of tokens from the diseases and genes collections with greater than 1,000 occurrences in the core corpus.

**Supplementary Figure 2.**
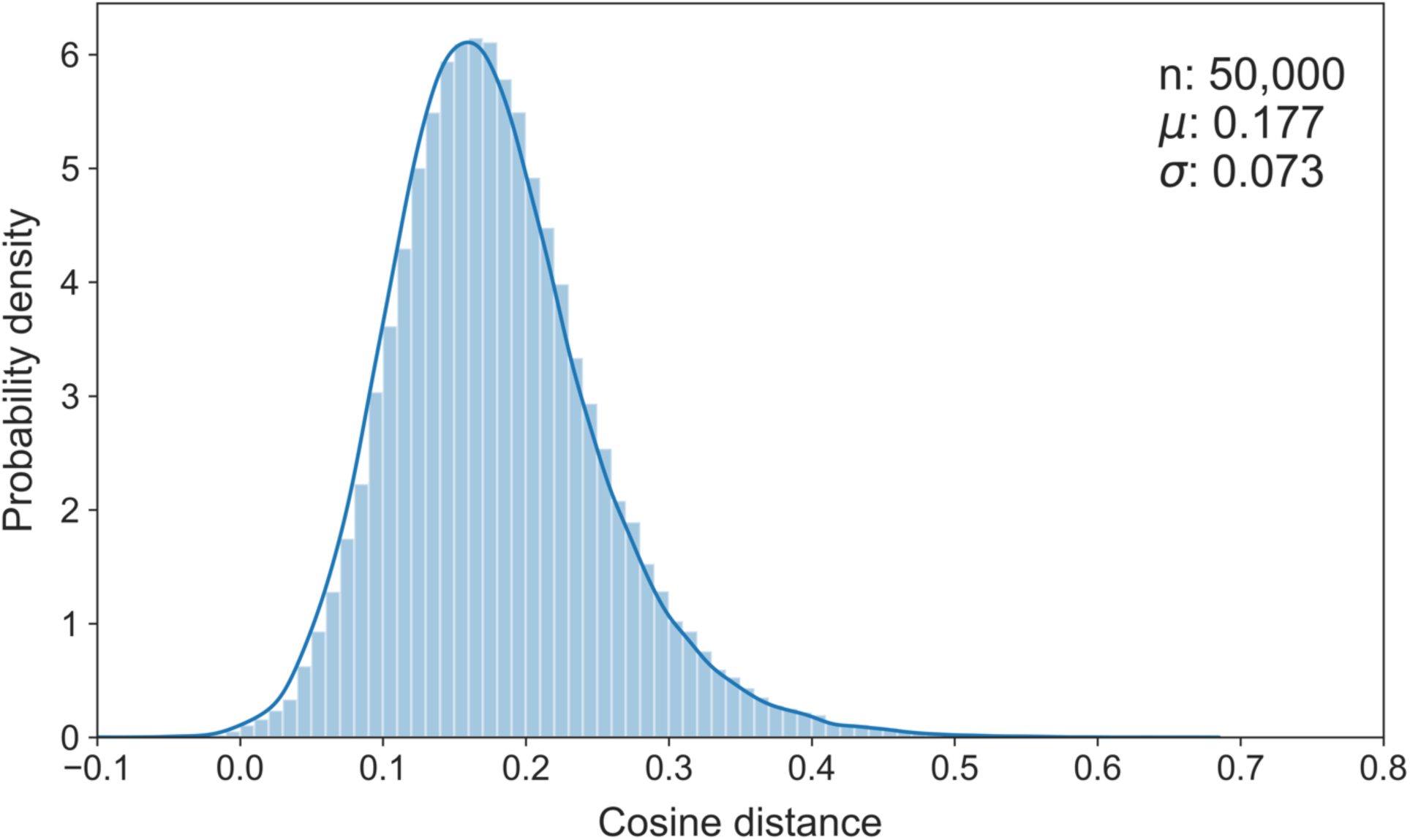
Distribution of cosine distances. Cosine distances were calculated from 50,000 pairs of tokens from the diseases and genes collections with greater than 1,000 occurrences in the core corpus.

**Supplementary Table 1.** Overview of the different corpora consumed for our approach, including corpora sources, and the number of documents, words, and vectors. Numbers are current as of February 27th, 2019.

| **Corpus** | **Example Sources** | **Documents** | **Words** | **Vectors** | **Temporal Support** |
| --- | --- | --- | --- | --- | --- |
| *Core Corpus* |  | *87,196,339* | *286,882,069,630* | *64,943,478* | *Yes\** |
| PubMed | -BioAssay<br>-GARD<br>-PubMed<br>-PubMed Central<br>-OMIM | 34,982,036 | 45,493,733,496 | 50,903,193 | Yes |
| Grants and Preprints | -arXiv<br>-bioRxiv<br>-ExPORTER<br>-Grants.gov<br>-NIH Grants | 3,235,834 | 1,478,493,947 | 3,648,362 | Yes |
| *Clinical Trials and SEC* |  | *1,077,020* | *13,848,226,551* | *9,636,935* | *Yes* |
| Clinical Trials | - ClinicalTrials.gov<br>-EU Clinical Trials Register<br>-Keynote Oncology Clinical Trials<br>-Novartis Clinical Trials<br>-YODA Project | 299,874 | 544,247,332 | 2,235,882 | Yes |
| SEC | -SEC | 777,146 | 13,302,714,355 | 8,003,482 | Yes |
| *Patents and Applications* |  | 10,963,950 | 158,615,007,383 | 95,814,123 | Yes |
| Patents Granted | -USPTO | 5,602,299 | 74,074,988,300 | 69,337,337 | Yes |
| Patent Applications | -USPTO | 5,361,651 | 81,263,779,169 | 84,970,621 | Yes |
| Media Corpus |  | 6,913,289 | 8,068,531,404 | 16,968,134 | Yes |
| Media Corpus Update |  | 6,913,289 | 10,617,245,065 | 19,130,863 | Yes |
| Lawsuits |  | 78,186 | 163,280,405 | 372,221 | Yes |
| Devices and Diagnostics |  | 8,634,787 | 3,714,427,372 | 5,601,960 | No |
| FDA and CDC |  | 1,791,401 | 6,081,923,060 | 6,589,551 | No |
| Wikipedia | -WikiDoc<br>-Wikipedia | 5,806,028 | 5,538,234,679 | 13,522,108 | No |
| Blogs and Conferences |  | 86,961 | 5,061,109 | 98,630 | No |
| BMIS |  | 1,723,317 | 4,438,521,136 | 7,486,777 | No |
| Patient Blogs |  | 494,918 | 481,572,244 | 2,133,862 | No |
| PDB | -PDB | 144,024 | 6,421,179,896 | 1,800,341 | No |
| DOAJ | -DOAJ | 3,277,935 | 903,640,377 | 5,232,759 | No |
| Companies |  | 899,289 | 1,749,120,647 | 4,670,100 | No |
| *Clinical Case Reports* |  | 35,057,325 | 567,604,939 | 2,973,878 | Yes |
| Clinical Case Reports (PubMed) | -PubMed<br>-PubMed Central | 34,982,036 | 421,383,772 | 2,597,601 | Yes |
| Clinical Case Reports (Others) |  | 75289 | 146,775,541 | 1,264,458 | Yes |
| Companies and Clinical Trials |  | 1,294,337 | 2,045,129,938 | 6,220,201 | Yes |
| Bookshelf | -dbGaP<br>-NCBI<br>Bookshelf | 104,431 | 566,811,454 | 3,102,885 | No |

**Supplementary Table 2.** Percentile conversion table for *LS* and *GS* in our approach. For the *X*th percentile, the corresponding *LS* is calculated as 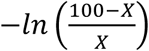 and the corresponding *GS* is calculated as 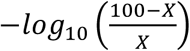.

| Percentile | *LS* | *GS* |
| --- | --- | --- |
| 99 | 4.61 | 2.00 |
| 95 | 3.00 | 1.30 |
| 90 | 2.30 | 1.00 |
| 85 | 1.90 | 0.82 |
| 80 | 1.61 | 0.70 |
| 75 | 1.39 | 0.60 |
| 50 | 0.69 | 0.30 |
| 25 | 0.29 | 0.13 |

## References

1 Landrum, M. J. et al. ClinVar: public archive of relationships among sequence variation and human phenotype. Nucleic Acids Res 42, D980–985, doi:10.1093/nar/gkt1113 (2014).

2 Stark, C. et al. BioGRID: a general repository for interaction datasets. Nucleic Acids Res 34, D535–539, doi:10.1093/nar/gkj109 (2006).

3 Tsuruoka, Y., Tsujii, J. & Ananiadou, S. FACTA: a text search engine for finding associated biomedical concepts. Bioinformatics 24, 2559–2560, doi:10.1093/bioinformatics/btn469 (2008).

4 Cheng, D. et al. PolySearch: a web-based text mining system for extracting relationships between human diseases, genes, mutations, drugs and metabolites. Nucleic Acids Res 36, W399–405, doi:10.1093/nar/gkn296 (2008).

5 Wei, C. H., Kao, H. Y. & Lu, Z. PubTator: a web-based text mining tool for assisting biocuration. Nucleic Acids Res 41, W518–522, doi:10.1093/nar/gkt441 (2013).

6 Mikolov, T., Sutskever, I., Chen, K., Corrado, G. & Dean, J. Distributed Representations of Words and Phrases and their Compositionality. arXiv e-prints (2013). <https://ui.adsabs.harvard.edu/\#abs/2013arXiv1310.4546M>.

7 Pennington, J., Socher, R. & Manning, C. in Proceedings of the 2014 Conference on Empirical Methods in Natural Language Processing (EMNLP). 1532–1543.

8 Joulin, A., Grave, E., Bojanowski, P. & Mikolov, T. Bag of Tricks for Efficient Text Classification. arXiv e-prints (2016). <https://ui.adsabs.harvard.edu/ \#abs/2016arXiv160701759J>.

9 Bojanowski, P., Grave, E., Joulin, A. & Mikolov, T. Enriching Word Vectors with Subword Information. arXiv e-prints (2016). <https://ui.adsabs.harvard.edu/\#abs/2016arXiv160704606B>.

10 Pyysalo, S., Ginter, F., Moen, H., Salakoski, T. & Ananiadou, S. Distributional semantics resources for biomedical text processing. Proceedings of LBM, 39–44 (2013).

11 Minarro-Gimenez, J. A., Marin-Alonso, O. & Samwald, M. Exploring the application of deep learning techniques on medical text corpora. Studies in health technology and informatics 205, 584–588 (2014).

12 Minarro-Gimenez, J. A., Marin-Alonso, O. & Samwald, M. Applying deep learning techniques on medical corpora from the World Wide Web: a prototypical system and evaluation. arXiv e-prints (2015). <https://ui.adsabs.harvard.edu/\#abs/2015arXiv150203682M>.

13 Bhasuran, B. & Natarajan, J. Automatic extraction of gene-disease associations from literature using joint ensemble learning. PLoS One 13, e0200699, doi:10.1371/journal.pone.0200699 (2018).

14 Cunningham, F. et al. Ensembl 2019. Nucleic Acids Res 47, D745–D751, doi:10.1093/nar/gky1113 (2019).

15 Hunt, S. E. et al. Ensembl variation resources. Database (Oxford) 2018, doi:10.1093/database/bay119 (2018).

16 Amberger, J., Bocchini, C. & Hamosh, A. A new face and new challenges for Online Mendelian Inheritance in Man (OMIM(R)). Hum Mutat 32, 564–567, doi:10.1002/humu.21466 (2011).

17 Amberger, J. S., Bocchini, C. A., Schiettecatte, F., Scott, A. F. & Hamosh, A. OMIM.org: Online Mendelian Inheritance in Man (OMIM(R)), an online catalog of human genes and genetic disorders. Nucleic Acids Res 43, D789–798, doi:10.1093/nar/gku1205 (2015).

18 Amberger, J. S. & Hamosh, A. Searching Online Mendelian Inheritance in Man (OMIM): A Knowledgebase of Human Genes and Genetic Phenotypes. Curr Protoc Bioinformatics 58, 1 2 1–1 2 12, doi:10.1002/cpbi.27 (2017).

19 Peters, M. E. et al. Deep contextualized word representations. arXiv e-prints (2018). <https://ui.adsabs.harvard.edu/abs/2018arXiv180205365P>.

20 Devlin, J., Chang, M.-W., Lee, K. & Toutanova, K. BERT: Pre-training of Deep Bidirectional Transformers for Language Understanding. arXiv e-prints (2018). <https://ui.adsabs.harvard.edu/abs/2018arXiv181004805D>.

21 Radford, A. et al. Language Models are Unsupervised Multitask Learners. (2019).

22 Nguyen, D. Q., Dras, M. & Johnson, M. A Novel Neural Network Model for Joint POS Tagging and Graph-based Dependency Parsing. arXiv e-prints (2017). <https://ui.adsabs.harvard.edu/\#abs/2017arXiv170505952N>.

23 Nguyen, D. Q. & Verspoor, K. An improved neural network model for joint POS tagging and dependency parsing. arXiv e-prints (2018). <https://ui.adsabs.harvard.edu/\#abs/2018arXiv180703955N>.

